# Reliance on polyfunctional tissue leads to a reproduction-immunity tradeoff due to inherent constraint

**DOI:** 10.1101/2021.01.28.428655

**Authors:** Vanika Gupta, Ashley M. Frank, Nick Matolka, Brian P. Lazzaro

## Abstract

The use of one tissue for multiple purposes can result in constraints, impaired function, and tradeoffs. The insect fat body performs remarkably diverse functions including metabolic control, reproductive provisioning, and systemic immune responses. Immunity and reproduction are observed to trade off in many organisms, although the mechanistic basis for the tradeoff is generally unknown. More generally, how do polyfunctional tissues simultaneously execute multiple distinct physiological functions? Using single-nucleus sequencing, we determined the *Drosophila melanogaster* fat body executes diverse basal functions with heterogenous cellular subpopulations. However, as an emergency function, the immune response engages the entire tissue. We found that reproductively active females exhibit impaired capacity to produce new protein in response to infection, resulting in the reproduction-immunity tradeoff. We suggest that such inherent internal limitations may provide a general explanation for the wide prevalence of physiological and evolutionary tradeoffs.

## Introduction

The need to balance multiple physiologically demanding and resource-intensive processes limits the ability of an organism to maximize performance in any one area. When two or more processes depend on a single tissue or resource pool, they unavoidably constrain each other, resulting in tradeoffs between the associated traits. Such tradeoffs are central to life history theory and affect the health, fitness, and evolution of all living organisms (1–3). Reproduction and immunity are two traits that trade off with each other across a broad diversity of systems (4,5) but the mechanisms and physiological constraints that underlie this tradeoff are poorly understood. In *Drosophila melanogaster* females, mating results in a rapid, endocrinologically-mediated drop in resistance to bacterial infection (6). We hypothesized that this tradeoff arises due to physiological constraints of using the same tissue, the abdominal fat body, for both reproductive investment and systemic immunity, and that understanding the basis for this tradeoff could serve as a model for understanding constraints on polyfunctional tissues in general.

The insect fat body is a highly multifunctional tissue that is engaged in central metabolic regulation, nutrient storage, detoxification of xenobiotics, reproductive egg provisioning, and mounting of systemic immune responses (7). Thus, this single tissue performs the functions of several vertebrate organs. The fat body is remarkably dynamic. For example, a bacterial infection significantly changes the expression of several hundred genes in the fat body of *Drosophila melanogaster*, including as much as 1000-fold induction of genes encoding antimicrobial peptides and marked down-regulation of glycolytic and basal metabolic pathways (8–10). Upon mating and sperm storage, the same tissue significantly upregulates genes involved in egg provisioning as the females increase their investment egg production (7). Reproduction and immune responses are both energetically demanding (11) and a female may need to simultaneously execute these processes as well as others. Given the finite number of cells and limited capacity for transcription and translation within each cell, how does one tissue achieve so many functions at once? Is the tissue composed of specialized subpopulations of cells that are individually devoted to each function? Or do all cells of the tissue perform all functions to a limited degree? When the tissue responds to stimulus, do the identities or sizes of cellular subpopulations change, or does each cell of the tissue alter its transcriptional profile in concert? Does the simultaneous execution of multiple processes by the single tissue constrain performance of each process?

To begin address these questions, we performed single-nucleus RNA sequencing (snRNAseq) on the fat bodies of *D. melanogaster* females in a replicated factorial design combining mating and bacterial infection. Mature adult female *D. melanogaster* were either mated in order to activate reproductive investment (M_) or held as virgin to limit reproductive investment (V_), and were either given a systemic bacterial infection with *Providencia rettgeri* to stimulate an immune response (_I) or were held uninfected (_U). We observed significantly lower survivorship of Mated-Infected (MI) females than Virgin-Infected (VI) females over 3 days post-infection (p = 0.0001; Fig. 1A), in accordance with previous observations (12,13) and demonstrating the expected tradeoff. We repeated each factorial treatment (VU, VI, MU, MI) in two independent biological replicates to generate a total of 8 samples for snRNAseq. From each sample, we dissected and pooled fat bodies from the abdomens of 40 female flies. The gut and ovaries are easily removed from the fat body tissue, but other cell types such as oenocytes, muscle cells, and hemocytes are harder to separate from fat body tissues and thus are co-isolated. We purified individual nuclei from the pooled tissues using a Dounce homogenizer followed by centrifugation onto a sucrose cushion (14). We performed snRNAseq using the 10X Genomics Chromium platform, loading at least 7000 nuclei per sample and sequencing at least 16,000 reads per nucleus for a minimum of 112 million reads per sample.

**Fig. 1.**
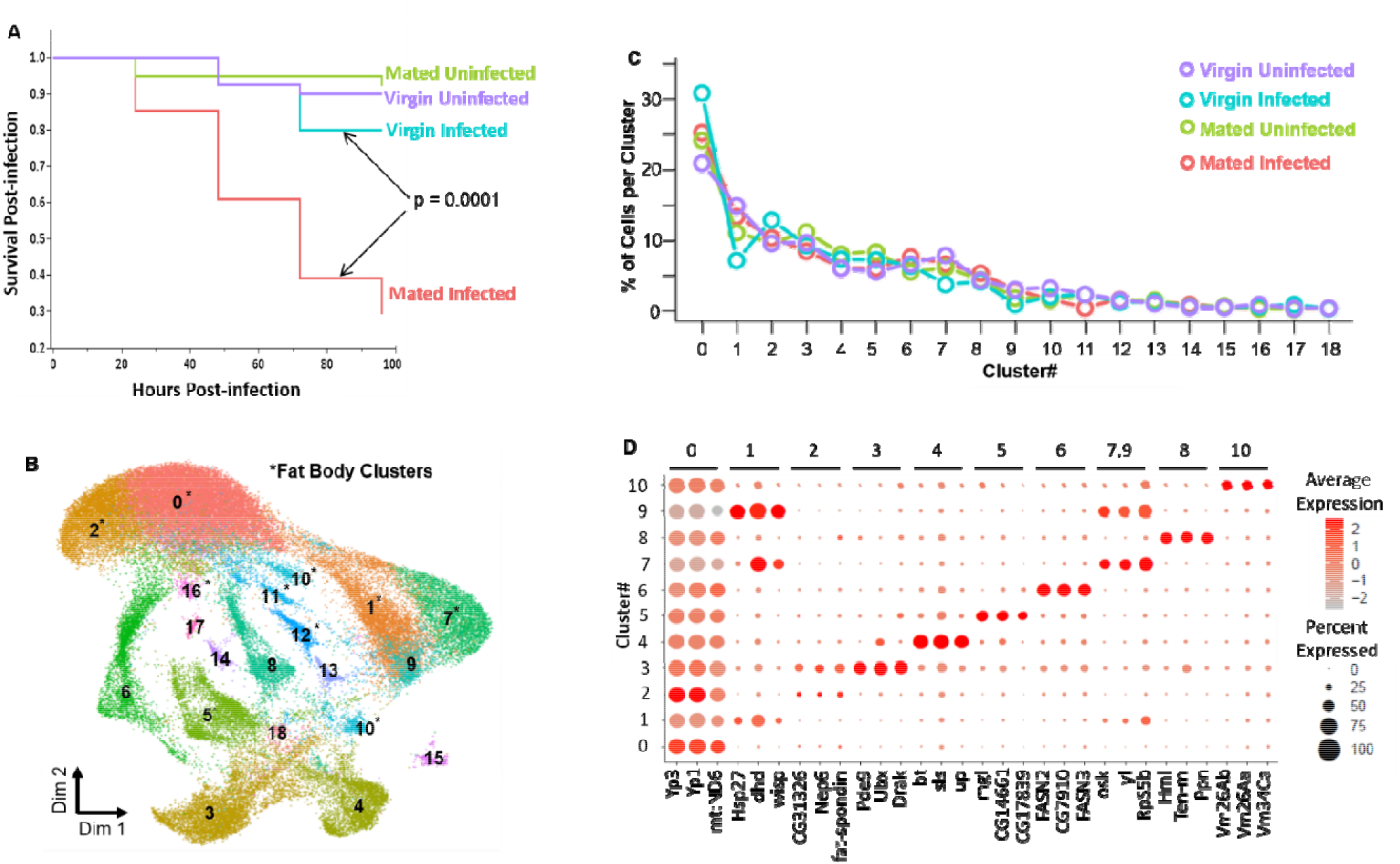
Single-nucleus sequencing of *Drosophila* fat body tissue. (A) Cox proportional hazard analysis showed that the mated *Drosophila melanogaster* females have significantly lower survival than virgin females (n =40; p = 0.0001) after infection with the Gram-negative bacterium *Providencia rettgeri*. Survival of uninfected virgin and mated females was not different over four days. (B) Combined Uniform Manifold Approximation and Projection (UMAP) of 56,000 nuclei from two replicates each of Virgin Uninfected, Virgin Infected, Mated Uninfected, and Mated Infected colored by their treatment identity. Clusters 0,1,2,5,7, and 10 marked with asterisk (*) represent subpopulations of the fat body tissue. (C) Percentage distribution of nuclei from four treatments (**<u>V</u>**irgin **<u>U</u>**ninfected, **<u>V</u>**irgin **<u>I</u>**nfected, **<u>M</u>**ated **<u>U</u>**ninfected, and **<u>M</u>**ated **<u>I</u>**nfected) across 19 clusters. All clusters are present in constant proportion across all four treatments. (D) Dot Plot showing expression of marker genes per cluster for top eleven clusters. Average expression of a marker gene in a cluster is represented by gradient of the colored dot and dot size represents the cell percentage per cluster expressing the marker.

We identified 19 expression clusters representing distinct cellular subpopulations (Fig. 1B, Fig. S1, S2) with 90% of the nuclei present in the eleven most abundant clusters (Fig. 1C). We assigned putative functional identities to each cluster based on the significantly high expression (p-adj <0.01) of diagnostic marker genes. Expression of top marker genes for the first eleven clusters is shown in Fig. 1D, and a full list of key expressed genes can be found in Table S1 and is illustrated in Fig. S3. The Supplementary Online Material contains detailed descriptions of the expression patterns and inferred functions for each cluster. We found significantly high expression of marker genes that are conventionally associated with fat body in six major clusters: 0, 1, 2, 5, 7 and 10. These six clusters contain approximately 60% of all the nuclei sequenced and demonstrate that the fat body tissue is composed of heterogeneous cell subtypes. An additional 5% of nuclei map to low-abundance fat body clusters (Clusters 11, 12, 13, and 16). Clusters 0 and 2 were defined by high expression of *yolk proteins 1* and *3* (Fig. 1D), while cluster 1, 5, 7 and 10 respectively had high expression of *deadhead, megalin, oskar*, and *vitelline membrane 26Ab*. All these marker genes are associated with oogenesis and egg development. Remarkably, the relative size of these clusters did not change significantly across the four treatment groups (Fig. 1C), indicating that the fat body does not respond to mating or infection by shifting the proportional representation of these specific cellular subpopulations. We infer that the remaining 35% of nuclei do not come from fat body tissue. Based on previously well-characterized cell-specific transcriptional markers, we determined that Cluster 4 is muscle (7% of sequenced nuclei), Cluster 6 is oenocytes (7%), Cluster 8 is hemocytes (5%), and Cluster 9 is uncharacterized (2%). Cluster 3 (10% of sequenced nuclei) remains uncharacterized but shows properties similar to both fat body and pericardial cells (see detailed description in Supplement, Fig. S4). These tissues are physically contiguous with the fat body, and interact with and have partially overlapping functions with the fat body tissue (15,16).

Upon mating, *D. melanogaster* females store sperm and begin to lay fertilized eggs, which requires increased investment in oogenesis (17). We asked whether the investment in reproduction varied across the six distinct subpopulations of the fat body tissue by cluster-specific differential gene expression analysis. When comparing virgin and mated females (24 hours post-mating) in the absence of infection, we found 186 differentially expressed genes across the six clusters with 145 genes significantly upregulated and 41 genes significantly downregulated (FDR <0.01; Table S2). We observed that none of the 186 genes were differentially regulated across all the six subpopulations (Fig. S5A) while 123 (66%) of these genes were differentially regulated in only one of the six subpopulations (Fig. S5A). For example, egg provisioning genes *yp1* and *yp3* were upregulated across four different clusters (Fig. S6A) while *yp2* was upregulated in only one cluster (Table S2). This indicates that the response to and investment in mating is heterogenous across fat body subpopulations. GO enrichment analysis of differentially regulated genes in each of the six subpopulations showed enrichment for diverse functions (Table S3). Upregulated genes in both Clusters 0 and 1 were enriched for one-carbon metabolism but mediated by two different mechanisms: s-adenosyl methionine (SAM; Cluster 0) and folate (Cluster 1). Cluster 1 also showed enriched upregulation of genes encoding ribosomal proteins, which were downregulated in Cluster 2. Upregulated genes in Cluster 2 showed enrichment for amino acid biosynthesis. We identified metabolic and detoxification pathways enriched in genes upregulated in Cluster 5, and upregulated genes in both Clusters 7 and 10 were related to phospholipase A1 activity. Therefore, while all six fat body subpopulations respond to mating stimulus, their heterogeneous response suggests subfunctionalization of the cellular populations.

The fat body mounts an intense and rapid immune response to bacterial infection (9,18) so we asked whether the whole tissue is engaged in that response or whether it maps to a restricted set of subpopulations. The answer, interestingly, is both. All clusters showed significant upregulation of immune response genes in both mated and virgin females, including genes that encode secreted antimicrobial peptides (Figs.S6B, S6C). However, the precise expression patterns were heterogeneous after infection, with particular combinations of immune genes induced most strongly in different subsets of clusters. Across the six major fat body subpopulations, 47 genes were induced by infection in both virgin and mated females. However, twice as many genes showed significant induction after infection in virgin females than in mated females (124 versus 63, FDR <0.01; Table S4, S5), indicating a negative impact of mating on the transcriptional response to infection. We found three genes (*attacin A, CG42807*, and *CG14322*) to be upregulated across all six fat body subpopulations in virgins (Table S4, Fig. S5B) while no genes were upregulated across every subpopulation in mated females (Table S5, Fig. S5C). Around 11% genes were upregulated in 4 or more of the six subpopulations in virgins compared to 6% in mated females. To understand the functional heterogeneity of the genes expressed in each cluster, we performed cluster-specific GO enrichment analysis of the genes that are differently expressed after infection in mated and virgin females separately (Table S6). Protein processing and secretion was a significantly enriched function of upregulated genes in Clusters 0 and 2 in both virgin and mated flies. Cluster 2 in virgin infected females also showed enrichment of genes for phagosome formation (Table S6). Downregulation of ribosome constituents was observed in Cluster 1 of mated flies and Cluster 2 of virgin females. We observed that 54% of differentially expressed genes in virgins and 65% of differentially expressed genes in mated were differentially regulated in only one of the six subpopulations (Figs.S5B, S5C). These data reveal heterogeneity in infection response across the fat body and demonstrate that the tissue-wide transcriptional response to infection is dampened by mating, probably contributing to the tradeoff between reproduction and immunity.

Most of the mating- and infection-induced transcriptional changes were heavily driven by clusters 0, 1, and 2 (Table S7), representing ~70% of all the nuclei from the six fat body subpopulations. We hypothesized that the involvement of such a large majority of fat body cells in resource-intensive physiological functions might constrain resource allocation, which could be reflected in coordinated regulation of gene expression networks or modules. To identify these modules, we constructed pseudotime trajectories from all the four treatments with Monocle (19–21), representing the transition of cells between differential functional states in response to mating or infection. An initial analysis revealed that the infected and uninfected fat body cells resolved into two completely disjointed trajectories defined by infection status. Trajectory 1 contained a majority of nuclei from VU and MU treatments while Trajectory 2 contained a majority of nuclei from VI and MI (Fig. 2A, Fig. S7). This suggests that fat body cells rapidly and dramatically change expression profile upon infection with no intermediate states visible at the 6-hour post-infection sampling time point. Only fat body cells (inferred using Seurat-based cluster analysis) were present in both of these trajectories (Fig. 2B). Other co-isolated cell types were present in only one of the two trajectories; indicating that they are not strongly transcriptionally responsive to infection. Using Louvain clustering in the two trajectories, we identified several modules of co-regulated genes that were enriched for specific functional ontologies. In Trajectory 1, we identified a module (Module 13, Fig. 2C, Table S8) with low aggregate expression score that was enriched in ribosome biogenesis (Fig. S8) in mated uninfected nuclei (MU) relative to virgin uninfected nuclei (VU), including CAP-dependent translation initiation factors. Surprisingly, the same set of genes (Module 16, Fig. 2C, Table S9) with the addition of one gene (*O-fucosyltransferase 2*) had a low aggregate expression score in mated infected (MI) nuclei contrasted to virgin infected nuclei (VI) in Trajectory 2 (Module 16). This suggests that protein synthesis might be reduced in the fat body after mating due to suppressed ribosome biogenesis. Furthermore, a subset of MI nuclei showed high expression of a module enriched in protein folding and degradation (Fig. S9), including genes involved in ER stress and unfolded protein response (UPR; Table S10). Electron microscopy confirmed dilated ER membranes in MI fat bodies (Fig. 3), indicative of ER stress (22). Since alleviation of ER stress is often attained via suppression of ribosome biogenesis to limit protein synthesis in the cell (22,23), a key factor underlying the observed reproduction-immunity tradeoff could be reduced capacity to produce immune-related proteins in mated females due to reduced protein synthesis.

**Fig. 2.**
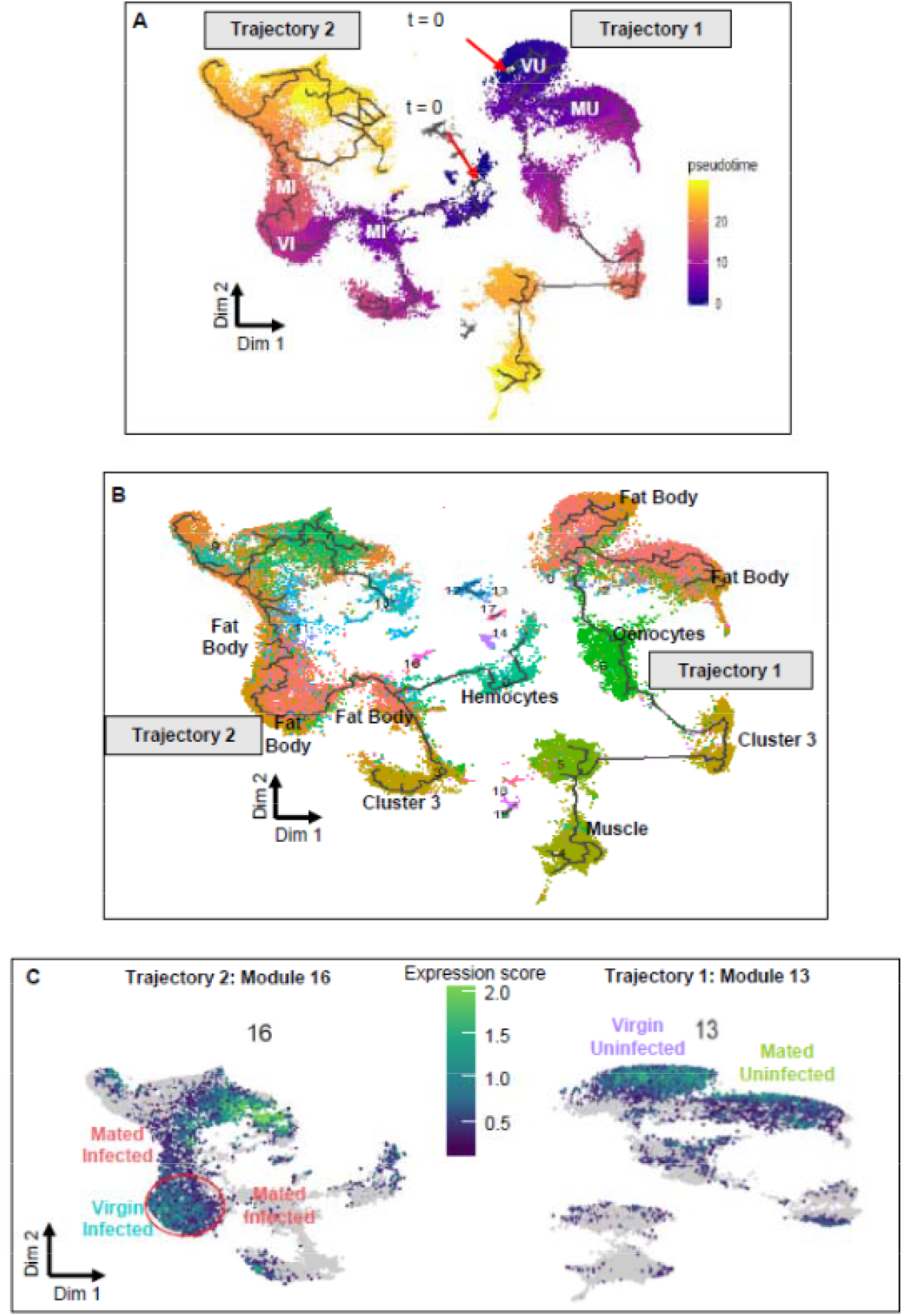
Pseudotime analysis showing differentially expressed gene modules. (A) Pseudotemporal ordering of nuclei along the two trajectories calculated from trajectory-specific (t = 0) points. Nuclei from four treatments (**<u>V</u>**irgin **<u>U</u>**ninfected (VU), **<u>V</u>**irgin **<u>I</u>**nfected (VI), **<u>M</u>**ated **<u>U</u>**ninfected (MU), and **<u>M</u>**ated **<u>I</u>**nfected (MI)) separate at different pseudo-time scales. The trajectories from infected nuclei are completely disjoint from the trajectories of uninfected nuclei, revealing a rapid and dramatic response to infection. (B) Monocle-based trajectory analysis separated nuclei along the two trajectories; colored by their cluster identity (from Figure 1B) show that only Fat Body nuclei are present in both trajectories. Other cell types such as oenocytes and muscle cells are present in Trajectory 1 and hemocytes are present in Trajectory 2, indicating these cell types do not have a strong transcriptional response to infection. (C) UMAP of Module 13 (Trajectory 1) and Module 16 (Trajectory 2) showing low gene aggregate expression scores for Mated Uninfected (Trajectory 1) and Mated Infected (Trajectory 2) compared to Virgin Uninfected and Virgin Infected respectively. Gradient of color represents the aggregate expression score with bright color indicating higher aggregate expression score. Each dot represents a single nucleus. GO term analysis showed enrichment for ribosome biogenesis in the two modules (Tables S8, S9, Fig. S8).

**Fig. 3.**
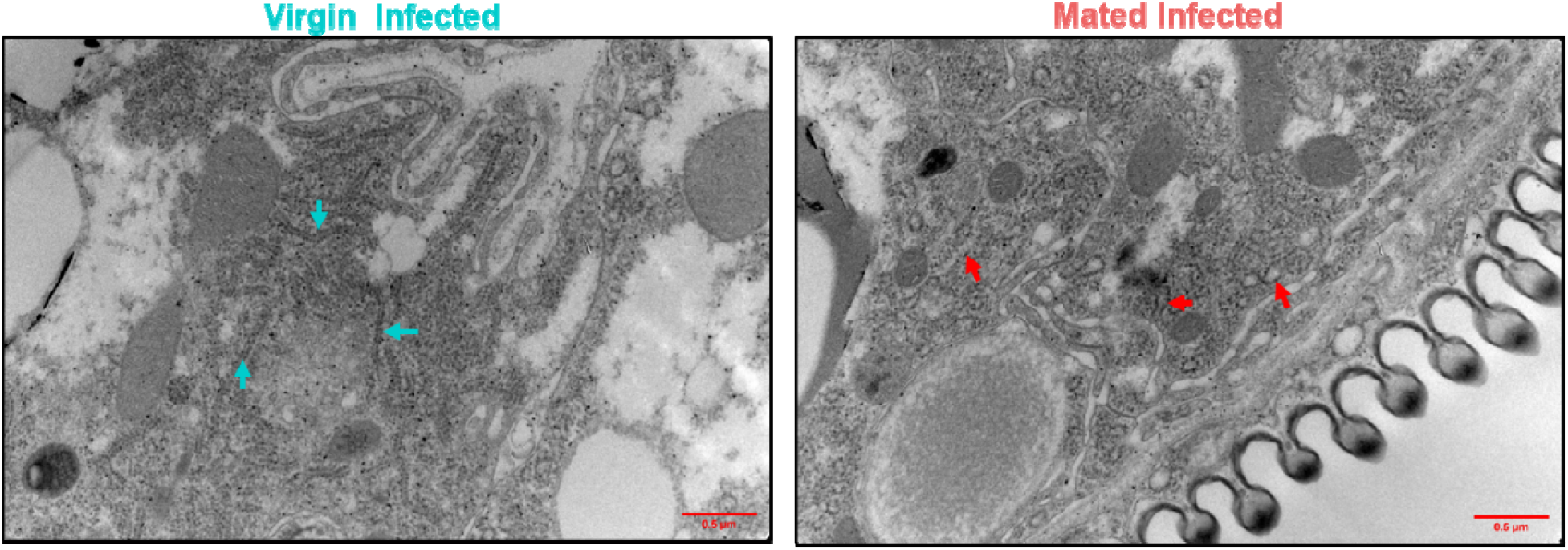
Electron micrographs of endoplasmic reticulum in the fat body (ER) Representative image (n = 4-5 images per treatment) showing dilation of ER membrane indicative of ER stress (right panel, red arrows) observed in fat body cells from Mated Infected females. Blue arrows show constricted ER membranes in Virgin Infected samples.

To test our hypothesis that mated infected (MI) females may lack sufficient capacity for translation in support of a full immune response to infection, we measured global protein synthesis in fat body tissues representing each of the four different treatments. We re-generated new female flies from each of the four factorial mating and infection treatments, dissected their fat bodies, and applied puromycin incorporation to label nascent polypeptides. Incorporated puromycin was then quantified on Western blots (20,21). We observed significant variability in global synthesis rates across the four treatments (one-way ANOVA, p=0.02, Table S11) with a spike in protein synthesis after infection in virgin females (VI) (Mean = 2.2, S.D. = 0.69) that fails to occur in mated (MI) females (Mean = 0.8, S.D. = 0.61) (Tukey’s HSD, p = 0.0005, Fig. 4A, 4B). These data are consistent with the reduction in ribosome biogenesis inferred from the sequencing data and with the hypothesis that the fat bodies of MI females are deficient in translation capacity. As the rapidity of an induced immune response is a critical determinant of infection outcome (22,23), a quantitative reduction or delay in the translation of immune response proteins such as antimicrobial peptides could contribute to the observed increased risk of death from infection in mated females.

**Fig. 4.**
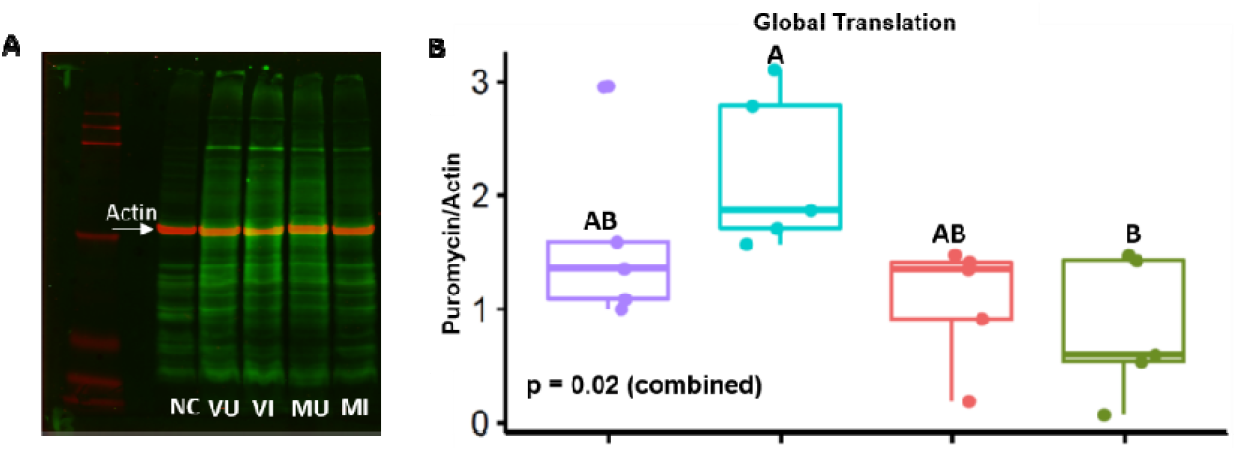
Effect of mating and infection on protein synthesis. (A) Representative image of puromycin incorporation in nascent polypeptides of the fat body tissue detected using anti-puromycin antibody and Western Blotting. Secondary antibodies labelled with different fluorophores detected puromycin (Green, 800nm) and Actin (Red, 700 nm). The fat bodies from Mated Infected (MI) produce noticeably less protein than those of the other treatments. Virgin Uninfected (VU), Virgin Infected (VI), Mated Uninfected (MU), and Mated Infected (MI) represent the four treatments. Negative Control (NC) (Figure A and D) shows proteins from fat body tissues which were not incubated with puromycin. (B) Quantification of relative protein synthesis using puromycin incorporation from four treatments (VU, VI, MU, and MI) (n = 5, 10 flies per treatment). Treatments not connected by same letter are significantly different (Tukey’s HSD, p < 0.005). Virgin Infected females synthesize significantly more protein than Mated Infected females. (C) Cox proportional hazard analysis showing rescued post-infection survival (p < 0.0001) of cycloheximide (CHX) pre-treated mated females (CHX-Mated Infected) compared to non-treated Mated Infected (n = 35-40 flies per treatment per replicate, three independent replicates). (D) Representative Western Blot image of puromycin incorporation from CHX pre-treatment of mated females (CHX). Results suggest CHX treatment partially rescues protein synthesis in response to infection compared to non-treated mated females (MI) (Paired Student’s t-test, p = 0.02). Puromycin incorporation was measured after six hours of infection.

Since we observed a reduction in protein synthesis in mated infected (MI) flies compared to all other groups, and especially compared to virgin infected (VI) females (Fig. 4B), we hypothesized that the high demand of producing reproduction-related proteins in mated females reduced capacity to translate new proteins in response to infection, even leading to ER stress in MI females. We predicted that the reproduction-immunity tradeoff could be alleviated if translational investment in reproductive proteins was reduced. To test this hypothesis, we mated females and then placed them on food containing cycloheximide (CHX) for 18 hours. CHX reversibly suppresses production of proteins in eukaryotes such as *Drosophila* (24). We subsequently transferred flies to food without CHX for six hours to allow them to clear the drug, and then gave them bacterial infections. Females that were treated with CHX after mating survived infection significantly better than mated females that were not treated with CHX (p<0.0001, Fig. 4C). We also observed an increase in post-infection protein synthesis in mated females pre-treated with CHX compared to non-treated females at six hours after infection (t (4) = 3.63, p=0.02, Fig. 4D, Table S12). Additional experiments confirmed that CHX has no direct role in the survival of infection (Fig. S10, see Supplemental Material for experimental details). Therefore, the observed tradeoff between reproduction and immunity is determined to be due to limited capacity for immune-related protein synthesis as a consequence of prior reproductive investment.

The impaired ability of the fat body to synthesize proteins in response to infection while simultaneously investing in reproduction illustrates a tradeoff driven by physiological constraint. The fact that immunity can be partially rescued with CHX treatment suggests the potential for plasticity in this tradeoff. Flies could, in theory, sustain greater immune capacity by reducing their commitment to reproductive investment. Thus, genetic variation for reproductive investment could allow evolutionary adaptation to infection risk. Reduced translation specifically in the fat body (25) extends lifespan in flies (26) through evolutionarily conserved mechanisms (27) observed across other organisms such as *C. elegans* (28,29) and mouse (30), whereas mating and reproduction are costly and negatively affects lifespan in fruit flies and other organisms (31). Translation in the fat body could therefore additionally be a mechanism mediating reproduction-longevity tradeoff. It seems likely that environmental factors, such as amino acid nutrition, may also influence the shape of these tradeoffs.

Managing competing physiological demands is a critical challenge for any polyfunctional tissue. We find here that the *Drosophila melanogaster* fat body executes diverse basal functions via heterogeneous cellular subpopulations. However, the whole tissue becomes engaged in an immune response. The gene expression markers that we have identified as defining the cellular subpopulations can serve to develop tags for future research into the dynamism and spatial structure of the *Drosophila* fat body. The fat body is a remarkable tissue that is highly responsive in regulating multiple aspects of physiology. However, while the fat body is enormously flexible, the shared reliance of multiple functions on a single tissue will inherently lead to constraints and tradeoffs. As we have shown in defining a reproduction-immunity tradeoff, compound stresses can overwhelm the tissue and lead to adverse outcomes. Understanding strategies that polyfunctional tissues use for balancing critical functions at the whole-tissue and sub-tissue levels can elucidate general mechanisms of physiological and evolutionary tradeoffs that underpin life history theory.

## Acknowledgments

We thank Peter Schweitzer for his assistance with sequencing. John Grazul and Katherine A. Spoth assisted with sample preparation and acquisition of electron microscopy images. We thank Profs. Mariana Wolfner, Robert Reed, Nicolas Buchon, and Nilay Yapici and Garrett League, Kathleen Gordon and Radhika Ravikumar for their feedback on the manuscript.

## Author contributions

Conceptualization: VG, BPL; Methodology: VG; Formal analysis: VG; Investigation: VG, AMF, NM; Writing: VG, BPL.

