## Supplement Text for "Reliance on polyfunctional tissue leads to a reproduction-immunity tradeoff due to inherent constraint"

Methods

**Fly Husbandry**

All experiments were performed using four-day post-eclosion *Drosophila melanogaster* females of the strain Canton S. Flies were raised on *ad libitum* cornmeal-sugar-yeast medium containing 6% Brewer’s yeast, 6% Cornmeal, 4% Sucrose, 0.7% Agar, 0.04% phosphoric acid and 0.004% propionic acid at 25°C on 12H:12H Light: Dark cycle.

**Mating treatment**

Females in this experiment are either virgin or mated. Virgins were collected by harvesting females within 6 hours of eclosion from the pupal case and housing them without males. To generate mated flies, 10 females were combined with 10 males in replicate vials for 24 hours prior to downstream experiments. The vials were observed for 30 minutes after males and females were combined to ensure that mating took place. More than 90% of females began mating within the first fifteen minutes. Males and females were held together for 24 hours to ensure that all the females mated.

**Infection treatment**

For the infection treatment, both virgin and mated females were infected with the Gram-negative bacterium *Providencia rettgeri*. Mated females were separated from males at the time of infection. To generate the bacterial inoculum, *P. rettgeri* were grown to saturation overnight in Lysogeny Broth (LB) broth at 37°C with shaking (200 rpm). Bacteria from the overnight culture were pelleted and resuspended in phosphate-buffered saline (PBS) to an optical density of A600 = 1.0. Flies were infected while under CO2 anesthesia by pricking the thorax with a 0.1 mm diameter Minutien pin that had been dipped in the bacterial suspension (1). This procedure delivers approximately 3000 bacteria to each fly. Uninfected females were handled in exactly the same way except they were not pricked with the needle.

**Post-infection survival assay**

To assay survival after infection, females were housed in groups of ten per vial, with the number of dead flies recorded every 24 hours for four days post-infection. Individuals surviving at the end of four days were censored. Post-infection survival analysis was done using Cox-Proportion Hazards model in JMP Pro v15. Pairwise risk ratios were calculated to find differences between treatments.

**Structure of the snRNA-seq experiment**

The snRNA-seq experiment was replicated twice across the four treatments: Virgin-Uninfected (VU), Virgin-Infected (VI), Mated-Uninfected (MU), and Mated-Infected (MI). All of the treatments were handled independently on separate days. The two replicate experiments were performed two months apart. A separate set of egg collections was done for each treatment. Flies were allowed to mature for four days prior to mating and infection treatments. Fat bodies of infected females were dissected at 6 hours post-infection.

**Fat body dissections and nuclei preparation**

Single-nuclei were isolated from abdominal fat body and associated tissues using the protocol described in Gupta and Lazzaro(1). Briefly, four-day old adults were anaesthetized using light CO_2_. The posterior tip of the abdomen was pulled out using fine forceps and the cuticle was cut laterally using spring scissors. The cuticle was carefully opened in cold adult hemolymph-like saline (HLS) and the gut and ovaries were removed. The cuticle and adherent issues were immediately transferred to chilled methanol fixative in order to preserve the quality of the RNA. After pooling tissues from 40 flies, cells were lysed in a hypotonic buffer using a Dounce homogenizer. Suspended nuclei were then purified using series of low-speed and sucrose gradient centrifugation steps. After purification, nuclei were suspended in PBS containing 2% BSA. If cell debris was observed, the nuclei suspension was filtered using a 20µm cell filter. Nuclei suspended in PBS (+2% BSA) on ice were immediately taken for 10X Chromium platform-based RNA sequencing. In pilot experiments, we found that this protocol dramatically improved sequencing quality and minimized mitochondrial contamination (2).

**10X Chromium nuclei preparation**

A small aliquot of the nuclei sample was incubated with Trypan Blue and the stained nuclei were counted using an automated cell counter. We used the total nuclei count to estimate sample volume required to sequence 7000 nuclei per sample on the 10X platform. RNA libraries were prepared using 10X chromium v3 chemistry and downstream sample processing was done as per the manufacturer’s protocol. Libraries were sequenced on Illumina platform following 10X recommended conditions.

**Data processing and analyses**

We used Cell Ranger (3) mkfastq (v3.0) (10x Genomics;) to generate demultiplexed FASTQ files from the raw sequencing reads. Reads were aligned to the Drosophila genome and quantified gene counts as UMIs using Cell Ranger count (v3.0) (10x Genomics). For snRNA-Seq reads, we counted reads mapping to introns as well as exons, as this results in a greater number of genes detected per nucleus, more nuclei passing quality control and better cell type identification, as previously described (4). To count introns during read mapping, we followed the approach described at <https://support.10xgenomics.com/single-cell-gene-expression/software/pipelines/latest/advanced/references>. Briefly, we built a ‘pre-mRNA’ Drosophila dmel-all-r6.24 reference using Cell Ranger (3) mkref (v3.0) (10x Genomics) with a modified gene transfer format (GTF) file Dmel v6 from Flybase. which was then used to align raw Illumina sequence reads to obtain sparse single cell expression matrix. The resulting matrices were analyzed using R (5) v3.5.3, and Seurat (6,7) v3.1. After performing sample QC using Seurat (6,7) and batch correction using Harmony (8) (Figure S1), we obtained data from 56,000 nuclei across all samples that were further processed to cluster nuclei into subpopulations (resolution 0.5) and identify cluster-specific canonical markers. Differential gene expression was evaluated using FindMarkers function in Seurat such that these genes were differentially expressed between treatments for both the replicates (FDR<1%) while not being differentially expressed between the two VU replicates.

We used Monocle (9–11) v3.0 to perform pseudotime analysis on the dataset. Gene modules containing co-varying genes were identified using stringency set to q-value<0.01. The gene sets obtained from these analyses were used for interpretation by performing Gene Ontology (GO) term enrichment analysis with g:Profiler (12) using default settings (p<0.01) for molecular function and biological process. GO: Profiler was also used to determine significantly enriched (p<0.01) KEGG pathways.

**Puromycin incorporation assay**

We measured global translation in the fat body using puromycin incorporation. Puromycin is an antibiotic which gets incorporated in nascent polypeptides during translation. Incorporated puromycin in polypeptides is then quantified using Western Blotting as per previously published protocols (13,14). The experiment was replicated five times. For each replicate of the experiment, ten adult female fat body tissues from each treatment were dissected in ice-cold adult hemolymph-like saline as described above. Tissues were stored on ice in 1 mL Schneider’s medium until ten tissues per sample were dissected and pooled. Tissues were carefully transferred to pre-warmed 1 ml Schneider’s medium containing 10µg/mL Puromycin. Samples were incubated at 25°C for one hour. After the incubation, 330µl of 50% trichloroacetic acid was added. Protein samples were prepared in SDS-PAGE loading buffer after washing tissues with 1M Tris base. Proteins were separated on 12% gel (Bio-Rad Mini-PROTEAN TGX Precast Protein Gels). Puromycin incorporation was assayed using anti-puromycin antibody (Millipore #12D10) and anti-actin antibody (Cell Signaling Technology #D6A8) was used to quantify actin as a loading control. Dual detection was perform using LiCor NIR fluorescent secondary antibodies - IRDye® 680RD Donkey-anti-Rabbit Antibody and IRDye® 800CW Donkey-anti-Mouse Antibody. Quantitative measurements were performed using LiCor Image Studio Lite and signal detected for each sample was relativized to its actin intensity. Difference in puromycin signal across treatments was determined using one-way ANOVA in JMP Pro v15 with “treatment” (VU, VI, MU, and MI) as the predictive factor.

**Cycloheximide feeding and survival assays**

We fed flies cycloheximide to inhibit translation and assay the effect of translation inhibition on immune defense in mated females. Flies were collected as virgins within six hours of eclosion and held in single-sex groups of ten flies per vial for 3-4 days. For the cycloheximide treatment, 50µl of 35 mM cycloheximide solution was put on the surface of food in vials and allowed to air dry. A dose of 35 mM has been previously used to inhibit translation in *D. melanogaster* (14–16). Virgin males and females were randomly chosen and three treatments were set up. In the virgin treatment (VI), females were held in single sex group of ten per vial in four replicate vials. Flies for the mating treatment (MI) were set up by combining 10 males with 10 females per vial in four replicate vials. For cycloheximide treatment (CHX), 10 males and 10 females were directly transferred to each of four replicate vials containing CHX-food. Flies were allowed to feed on CHX for 18 hours; after which they were transferred to fresh vials with no CHX to allow flies to recover and clear CHX. Flies recovering within 6-12 hours of cycloheximide treatment has been shown in (17). Six to seven hours later, flies were infected with *P*. *rettgeri* as described above. Three independent replicate blocks were generated following this protocol. The effect of cycloheximide on post-infection translation was assayed using puromycin incorporation and Western blotting as described above. To test for difference in translation between CHX and MI, a paired t-test was done in R (5) v3.5.3. Box plots were plotted using package ggpubr (15) in R v3.5.3.

Our experimental results showed that CHX treatment of mated females improved their ability to survive infection. To validate that immunity was enhanced in mated females by CHX-inhibition of translation only when CHX was delivered after mating but before infection, we performed additional control experiments that included three treatments in addition to the VI, MI, and CHX treatments described above. In the first of these, virgin females were transferred to CHX-containing (CHX-V) food for 18 hours, after which they were transferred to fresh vials with no CHX for six hours prior to infection. Four such vials were set up with ten flies per vial. This treatment mirrors the MI treatment, except the flies are not mated, and it tests whether CHX has a directly protective effect against infection. For the second treatment, flies were fed with CHX **c**ontinuously (CHX-C). In this treatment, females were transferred to CHX-treated food at the time of mating and were maintained on CHX food for the duration of the experiment, including after infection. For the third treatment, females were combined with males on normal food and these flies were transferred on CHX food only post infection with bacteria (CHX-PI). Thus, in this treatment, the females will be fully invested in reproduction and will have translation impaired as they initiate their immune response. These additional treatments were replicated three times with post-infection mortality in all experiments recorded every 24 hours for four days. The individuals surviving at the end of four days were censored in the analysis. The survivorship data are shown in Fig S10. We found that inhibiting translation prior to mating but returning the flies to food without CHX prior to infection (the original CHX treatment) significantly improves survivorship of infection (p < 0.0001) relative to females who are not provided with CHX in the food (MI). However, if CHX is provided continuously (CHX-C) or after infection (CHX-PI), survivorship is the same or worse as is observed for MI flies and is significantly worse in than is observed for VI, CHX-VI, or CHX flies (p < 0.0001). There was no difference in survivorship between VI and CHX-VI flies (p =0.69). Thus, CHX treatment only promotes survivorship of infection when the CHX limits reproductive investment without impairing immune-related translation.

**Electron microscopy**

### Following the protocol described for mating and infection as described above, flies for four different treatments (VU, VI, MU, MI) were set-up and 5-10 tissues per treatment were dissected. Tissues were immediately fixed in 2% glutaraldehyde, 2.5% formaldehyde, 0.1 M Na cacodylate, pH 7.4., and then transferred into 2.5% glutaraldehyde in 0.05M cacodylate buffer pH 7.4 for 2 hours at 4°C. Samples were post-fixed in 1% OsO_4,_ 0.05M cacodylate buffer for 1 hour at 4°C. The samples were dehydrated in 25% ethanol (4°C for 15’) and 50% ETOH (4°C for 15’). We placed the samples in 2% uranyl acetate in 70% ETOH at 4°C for 48 hours. After 48 hours, serial dehydration was conducted in ethanol (first 95% then100%) followed by two washes with 100% acetone. Both the dehydration and washing steps lasted 10 minutes each and were done at 4 degrees Celsius. The abdomens were infiltrated with Embed 812 (EMS #14120). The abdomens were then embedded in flat molds and polymerized at 60 degrees Celsius for 24 hours. The samples were ultramicrotomed on a Leica Ultracut UTC 7 using a Diatome 6 degree knife. 60-70nm thick sections were mounted on 50 mesh copper grids coated with polyvinyl Butvar/carbon grids. Grids were stained for 15 minutes with 2% uranyl acetate and rinsed with water. The imaging was done with a 120 Thermo-Fisher Tecnai T12 BioTwin at 1200kV. Images were obtained using a Gatan 794 CCD camera.

Supplementary Text

***Cluster Descriptions***

We performed single-nucleus RNA-sequencing on 56,000 nuclei from eight samples (two replicates each of VU, VI, MU and MI treatments). After confirming sequence quality, we performed batch correction using Harmony with default parameters (Figs.S1,2), and clustering using Louvain algorithm with resolution set to 0.5 and default parameters using Seurat (v3.1). This yielded 19 distinct nuclear (cell) subpopulations. The subpopulations are numbered from 0 to 18 in descending order of size, the largest being “Cluster 0” and the smallest being “Cluster 18”.

We performed differential gene expression analysis for each cluster against all other clusters using the function FindConservedMarkers in Seurat to identify genes that were significantly upregulated (FDR <1%) or overexpressed in each cluster across all the four treatments (VU, VI, MU, and MI) when compared to nuclei in all other clusters (Fig.S3, Table S1). These significantly upregulated genes are also called “canonical markers” or “marker genes” or “markers” for each cluster. Next, we performed GO (Gene Ontology) term enrichment (p<0.01) analysis on each list of “marker genes” corresponding to each cluster using g:Profiler (12)**.** We inferred the function sub-type of each of the clusters based on enrichment of GO terms associated biological processes, molecular function, and cellular component along with KEGG pathway enrichment. Some clusters, as expected, appear to belong to cells of other tissues that were co-dissected with the fat body. These expressed distinctive marker genes diagnostic of cell types such as muscle, oenocytes, hemocytes, and crystal cells. We annotated the remaining clusters as fat body subtypes. Below we discuss each cluster in terms of its size (in terms of fraction of total nuclei in the tissue), key marker genes expressed (Supplementary Table S13), and significant GO terms (Supplementary Table S14), and we summarize these findings by inferring a putative function for each cluster.

**Cluster 0 – Fat body cells**

Cluster 0 is the primary cluster containing fat body cells and contains 25% of all the nuclei sequenced. This is the largest and most generic cluster of fat body cells and it has the fewest distinctively expressed marker genes, with only 7 diagnostic marker genes (Table S13). The genes *yolk protein* 3 (*yp3*) and *yolk protein* (*yp1)* are the top two markers expressed in the cluster. Yolk proteins in *Drosophila* are expressed and secreted by the fat body for their uptake by mature oocytes (16). Additionally, we also found that *Phosphoglucose mutase 1* (*Pgm1*), an enzyme involved in glycolysis, was one of the marker genes. Enriched GO terms included phospholipase A1 activity (Table S14). We conclude that the primary roles of Cluster 0 are producing yolk proteins and metabolism.

**Cluster 1 – Fat body cells**

Cluster 1 consists of 12% of all nuclei sequenced and is defined by 77 markers (Table S 13). The large number of marker genes indicates that this cluster may be a specialized sub-type of the fat body tissue. *Heat shock protein 27 (Hsp27)* and *deadhead (dhd)* are the top markers expressed in this cluster. Enriched GO terms included RNA binding, organelle organization, regulation of metabolic and cellular processes (Table S14). The modENCODE bulk RNAseq dataset represented in FlyBase ([www.flybase.org](http://www.flybase.org)) indicates that nine out of the top ten marker genes showed moderately high (minimum RPKM = 29) to very high expression (minimum RPKM = 110) in four-day old carcass. Interestingly, the modENCODE bulk RNAseq dataset showed that these genes are also expressed in the in ovaries, although the ovaries were removed in our tissue dissections. Fat body Cluster 1 may have a role in reproduction.

**Cluster 2 – Fat body cells**

Cluster 2 consists of 11% of nuclei sequenced and was marked by 23 genes (Table S13). The genes *yp3*, *yp1*, and *trehalose-6-phosphate synthase 1* (*Tps1)* were the markers with the highest expression. The top markers of cluster 0 (*yp1*, *yp3* and *Pgm1*) also marked cluster 2. The GO enrichment term “lipase activity” characterized Cluster 2 (Table S14) and KEGG pathways for starch, sucrose, and galactose metabolism were significantly enriched in expression (Table S14). This suggests that, like Cluster 0, cluster 2 is a fat body cluster that has metabolic functions and is involved in reproductive provisioning.

**Cluster 3 – Unknown cell type**

Cluster 3 consists of 10% of sequenced nuclei and distinctively expressed 284 marker genes (Table S13). Top markers of cluster 3 are *Epidermal growth factor receptor* (*Egfr*), *nicotinic acetylcholine Receptor α7* (*nARCH7α*), and *Death-associated protein kinase related (Drak*). Cluster 3 was also marked by expression of genes in the Toll (*Dif*, *Tl*), *Imd* and JAK/STAT (*Stat92E*), and insulin signaling (*thor*, *foxo*) which are immune-related and metabolic pathways. Cluster 3 also expressed as a marker gene *adiponectin receptor* (*AdipoR*), which controls insulin signaling to regulate metabolism and maintain germ-line stem cell populations. Enriched GO terms included biological processes such as response to stimulus and positive regulation of biological process (Table S14). KEGG pathway analysis showed enrichment for purine metabolism (Table S14). Availability of *nARCH7α-*GAL4 driver (17) allowed us to spatially resolve cluster 3 (Fig.S4). Surprisingly, expression of the fluorescent reporter mCherry under the control of nARCH*7a* driver coincided with the location and morphology of pericardial cells (18,19). Therefore, although this cluster appears to be involved in immunity and metabolism and has expression patterns indicative of a fat body subpopulation, the physical location and morphology suggest pericardial cells. In the absence of definitive certainty, we conservatively consider this cell type to have an unknown tissue identity.

**Cluster 4 – Muscle cells**

Cluster 4 consists of 7% of all nuclei sequenced and is defined by 277 marker genes (Table S13). The top two most expressed genes of the cluster are *bent (bt),* and *sallimus (Sls)*. Enriched GO terms included biological processes related to striated muscle cell development, muscle cell differentiation, and actomyosin structure organization (Table S14). Therefore, we marked cluster 4 as muscle cells.

**Cluster 5 – Proliferative fat body cells**

Cluster 5 consists of 7% of the sequenced nuclei and was marked by expression of 184 genes (Table S13). Genes *megalin (mgl)* and *CG14661* were the top markers of this cluster. *Megalin* is a glycoprotein and regulates endocytosis (20). Marker gene *grainyhead* expressed in cluster 5 is a transcription factor of FGFR signaling pathway which promotes wound healing (21), cell proliferation, and cell growth; *grainyhead* has been previously implicated in determining tolerance of bacterial infection (22). Transcription factor *vvl* was a marker for this Cluster 5 that has been previously shown to upregulate expression of the antimicrobial peptide gene *cecropin* expression in the fat body (23). Enriched GO terms included (Table S14) biological processes associated with multicellular organism development. The hippo signaling pathway, as well as phenylalanine and tyrosine metabolism were significantly enriched KEGG pathways (Table S14). Expression of *grainyhead* and genes in the hippo pathway suggests that the cells in Cluster 5 may be proliferative (24).

**Cluster 6 – Oenocytes**

Cluster 6 consisted of 7% of nuclei and expressed 124 marker genes (Table S13). The top genes marking the cluster are *FASN2* and *CG7910*. When comparing the markers of this cluster with previously published gene lists enriched in oenocytes and fat body, we found that 37 out of 124 marker genes overlapped with previously published oenocyte-expressed genes (25). Enriched GO terms included biological processes related to fatty acid metabolic process (Table S14). Since oenocytes in *Drosophila* are primarily responsible for fatty acid biosynthesis (25) and there is high overlap between this cluster and previously defined oenocyte markers, we conclude cluster 6 nuclei are from oenocytes.

**Cluster 7 – Reproductive provisioning fat body cells**

Cluster 7 contained 6% of the nuclei sequenced. Cluster 7 was marked by expression of 617 genes (Table S13). *Oskar (Osk)*, *deadhead (dhd*), and *yolkless* (*yl*) are the top marker genes of cluster 7. Enriched GO terms included enrichment of the biological processes’ terms cellular process, cellular component organization, sexual reproduction and many others (Table S14). Enriched KEGG pathways included phagosome and mTOR signaling pathway. These expression patterns indicate that that cluster 7 is a fat body subpopulation whose functions include reproduction.

**Cluster 8 – Hemocytes**

Cluster 8 was formed by 5% of the sequenced nuclei and was marked by expression of 208 marker genes (Table S13) including *hemolectin (*hml), *tenascin major (Ten-m)*, and *papillin (ppn)*. Enriched GO terms included biological processes related to anatomical structure and animal organ morphogenesis, and enriched KEGG pathways included ECM-receptor interaction, MAPK signaling pathway, and lysosome activity (Table S14). *Hemolectin* is a diagnostic marker (26) of hemocytes and lysosomal activity is a well-characterized phagocyte function so we labelled cluster 8 as hemocytes. Hemocytes have previously been shown to be in close physical association with the fat body, where they act as sentinels of infection and promote expression of immune response genes by the fat body (27).

**Cluster 9 – Uncharacterized**

Cluster 9 consisted of 2% of sequenced nuclei and expressed 877 marker genes (Table S13). *Hsp27* and *wispy (wsp*) are the top two markers of cluster 9. GO enrichment analysis showed that about 400 genes out of 877 were enriched for biological process related to biogenesis and regulation of cellular processes (Table S14). 227 genes out of 877 genes were enriched for reproduction (Table S14). KEGG pathways enriched are Ubiquitin-mediated proteolysis, NOTCH signaling, RNA transport and spliceosome (Table S14). However, we could not interpret any definitive functional pattern. Cluster 9 therefore remains uncharacterized in our dataset.

**Cluster 10 – Structural fat body cells**

Cluster 10 consists of 2% of sequenced nuclei and expresses 54 marker genes (Table S13). Top markers of the cluster included vitelline membrane proteins (*Vm24Aa*, *Vm34Ac*), and *trol*. Laminins (*Lan-A*), perlecan (*trol*) along with heparan sulphate proteoglycan (HSPG) were also markers for this cluster. Together, these genes form CIVICs (Collagen IV Intercellular Concentrations) in the basement membrane of the fat body (28). CIVICs are a key to inter-adipocyte adhesion in the fat body (28). Enriched GO terms included anatomical structure development, cell development and differentiation (Table S14). Extra-Cellular Matrix and ECM-receptor interaction was significantly enriched as a KEGG pathway. We concluded that cluster 10 plays a role in maintaining structure of the fat body.

**Cluster 11 – Chorion producing fat body cells**

Cluster 11 consists of 1.9% nuclei of total nuclei. Ten genes marked the cluster 11, seven of which express chorion proteins (Table S13). The top markers of cluster 11 are *Chorion protein 36 (Cp36*) and *Chorion protein 38 (Cp38*). Enriched GO terms included biological process: chorion-containing eggshell formation (Table S14). Dobson et al (29) (2016) reported expression of chorion proteins in tissues other than ovaries. This cluster probably represents a subpopulation of fat body tissue producing chorion proteins for transport to developing oocytes.

**Cluster 12 – Catabolic fat body cells**

Cluster 12 consists of 1.5% of the sequenced nuclei and was marked by 55 expressed genes (Table S13). *Mucin-related-18B* (*Mur18B*) and *CG14292* are the top markers of cluster 12. Enriched GO terms included biological processes associated with ion transport (Table S14). Enriched KEGG pathways were phagosome, oxidative phosphorylation, and metabolic pathways (Table S14). Several genes expressed in these cells encoded vacuolar ATPases (V-ATPases) which have functions in phagocytosis, lysosomal activity, and participate in the mTOR pathway. In larval fat body cells, V-ATPases has previously been shown to play a role in lysosomal formation, acidification and cargo degradation in larval fat body (30). These cells could be playing a similar role in in the adult fat body as well.

**Cluster 13 – Putative digestive fat body cells**

Cluster 13 consisted of 1% of sequence nuclei and was marked by expression of nine genes (Table S13). *α-Trypsin (αTry)* and *Jonah 65Aiv (jon65Aiv)* are the top marker genes expressed in this cluster. Enriched GO terms included proteolysis biological process. This cluster also expressed three Jonah proteins (Table S14). Although Jonah proteins are highly expressed in the adult midgut (31), their expression has also been reported in larval fat body (32,33) and fly head (34). A similar cluster marked by *αTry* and Jonah proteins was previously reported in single nucleus sequencing of male fat bodies and was termed as a putative digestive cell cluster (35). We retain that notation for cluster 13.

**Cluster 14 – Crystal cells**

Cluster 14 consists of 0.7% of the sequenced nuclei and was marked by expression of 79 genes (Table S13). The top two markers of cluster 14 are *prophenoloxidase 1 (PPO1*) and *prophenoloxidase 2* (*PPO2*)*.* Enriched GO terms included embryo, reproductive structure, and reproductive system development (Table S14). ECM-receptor interaction was the enriched KEGG pathway (Table S14). Given that *PPO1 and PPO2* are predominantly expressed in crystal cells (36), a comparatively rare hemocyte type, we infer cluster 14 to be crystal cells.

**Cluster 15 – Neuronal**

Cluster 15 consists of 0.6% of sequenced nuclei and is marked by expression of 168 marker genes (Table S13). Highly expressed markers of the clusters include *paralytic (para)*, *shaker*, *resistant to dieldrin* (*rdl*). GO term analysis showed enrichment of biological processes such as anterograde trans-synaptic signaling, synaptic signaling, and nervous system development (Table S14). The predominant function associated with Cluster 15 was neurotransmission and we conclude that this cluster represents neuronal cells.

**Cluster 16 – Stress-response fat body cells**

Cluster 16 consists of 0.6% of sequenced nuclei and was marked by four transmembrane proteins: *what else (whe*), *la costa (lcs*), *CG45080, and CG16826* (Table S13). We did not find enrichment for GO terms using the four marker genes of this cluster (Table S14). *Lcs* expression is reported in larval fat body (37). *Whe* and *lcs* mediate Cyclin J-dependent gut recovery upon bacterial digestion in fruit flies (38). Therefore, this cluster may play an important role in alleviating stress response possibly including bacterial infection.

**Cluster 17 – Nephrocytes**

Cluster 17 consists of 0.5% of sequenced nuclei (Table S13). The cluster showed expression of 38 marker genes. *Cubilin* and *CG42255* are the top marker genes. GO term analysis (Table S14) showed that the genes expressed in this cluster were enriched for biological processes in nephrocyte diaphragm assembly and nephrocyte filtration, suggesting that this cluster represents nephrocytes.

**Cluster 18 – Tracheal cells**

Cluster 18 consists of 0.4% of sequenced nuclei with 122 genes marking cluster 18 (Table S13). The top two markers expressed in cluster 18 are *antennapedia* (*Antp*) and *waterproof (wat)*. GO term analysis showed expression of *Waterproof* along with 17 other genes (Table S14) enriched for biological process open tracheal development. This suggests Cluster 18 to be tracheal cells.

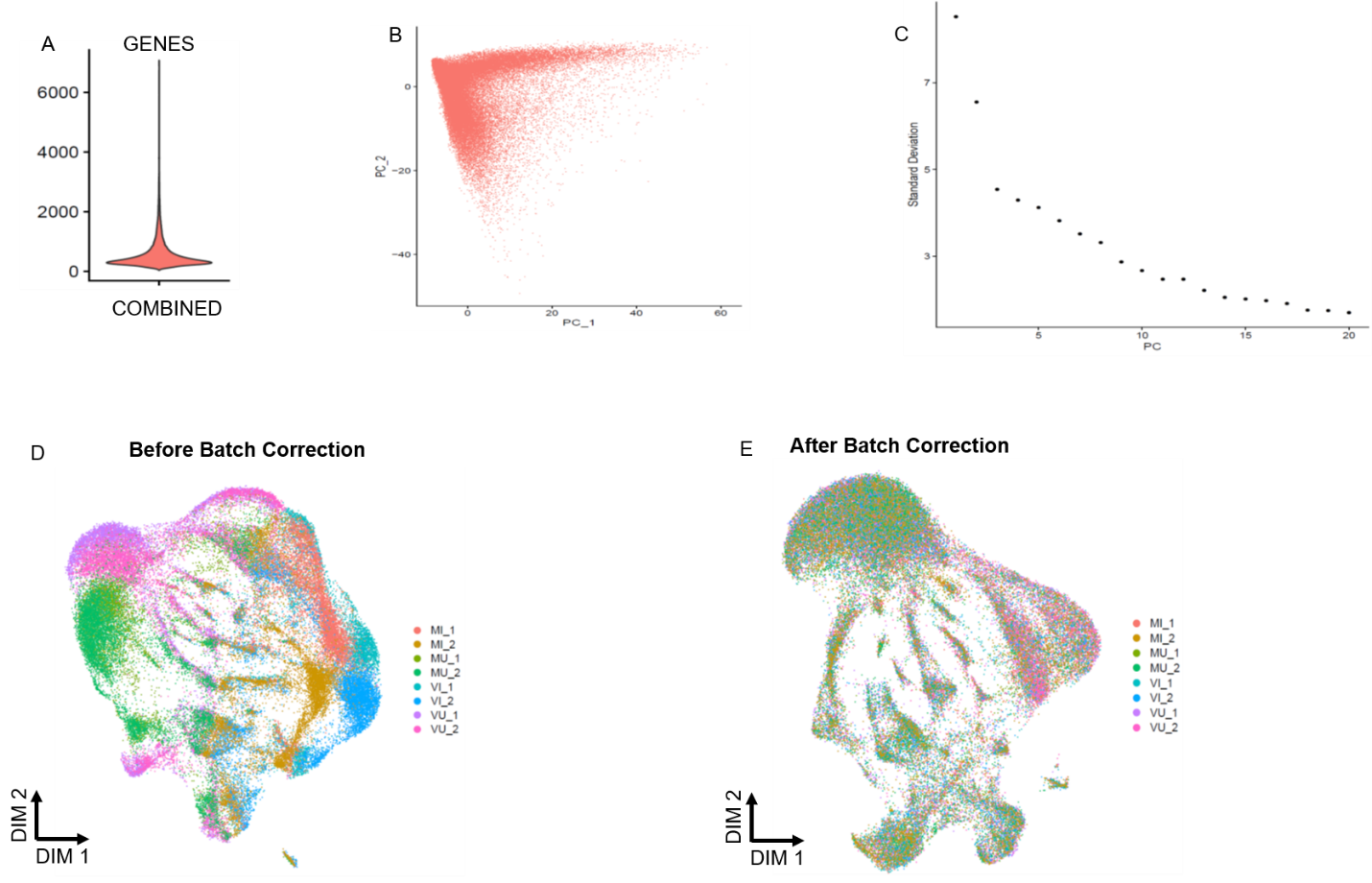

**Fig.S1. Quality Control of single-nuclei sequencing data of fat body cells of *Drosophila melanogaster*.**

(A) Distribution of the number of genes detected per sequenced nucleus across the entire experiment. (B) PCA plot of gene expression data showing first two principal components showing that there are no clear outliers in the dataset. (C) Elbow plot ranking variance explained by each principal component. (D) Uniform Manifold Approximation and Projection (UMAP) of nuclei from all the samples before batch correction. Nuclei from each replicated treatment are labeled with different colors (Mated Infected (MI), Mated Uninfected (MU), Virgin Infected (VI), and Virgin Uninfected (VU)). (E) UMAP of nuclei from the entire dataset after Harmony-based batch correction. After performing sample QC using Seurat (6,7) and batch correction using Harmony (8), we obtained data from 56,000 nuclei across all samples with a median of 399 genes and 851 RNA molecules per nucleus.

Number of Genes per cluster

Total No. of RNA molecules per cluster

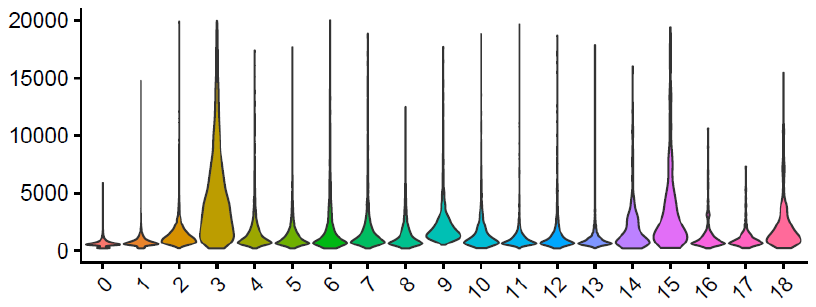

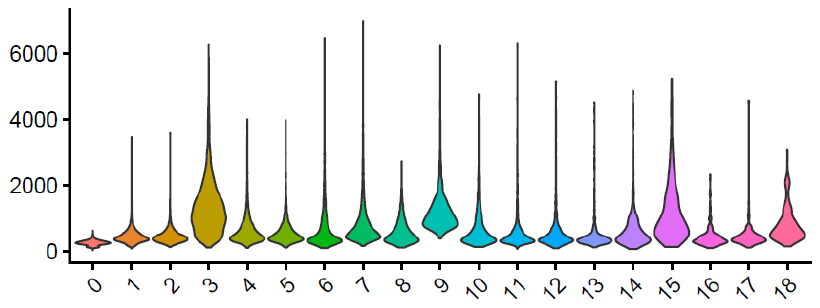

**Cluster#**

A

B

**Fig.S2. Metrics of single-nuclei sequencing data from *D. melanogaster* fat body.**

(A) Number of RNA molecules detected per cluster. Clusters (x-axis) represent 56,000 nuclei from eight samples. (B) Number of unique genes identified per cluster. The plot shows clusters (x-axis) from 56,000 nuclei sequenced across eight samples.

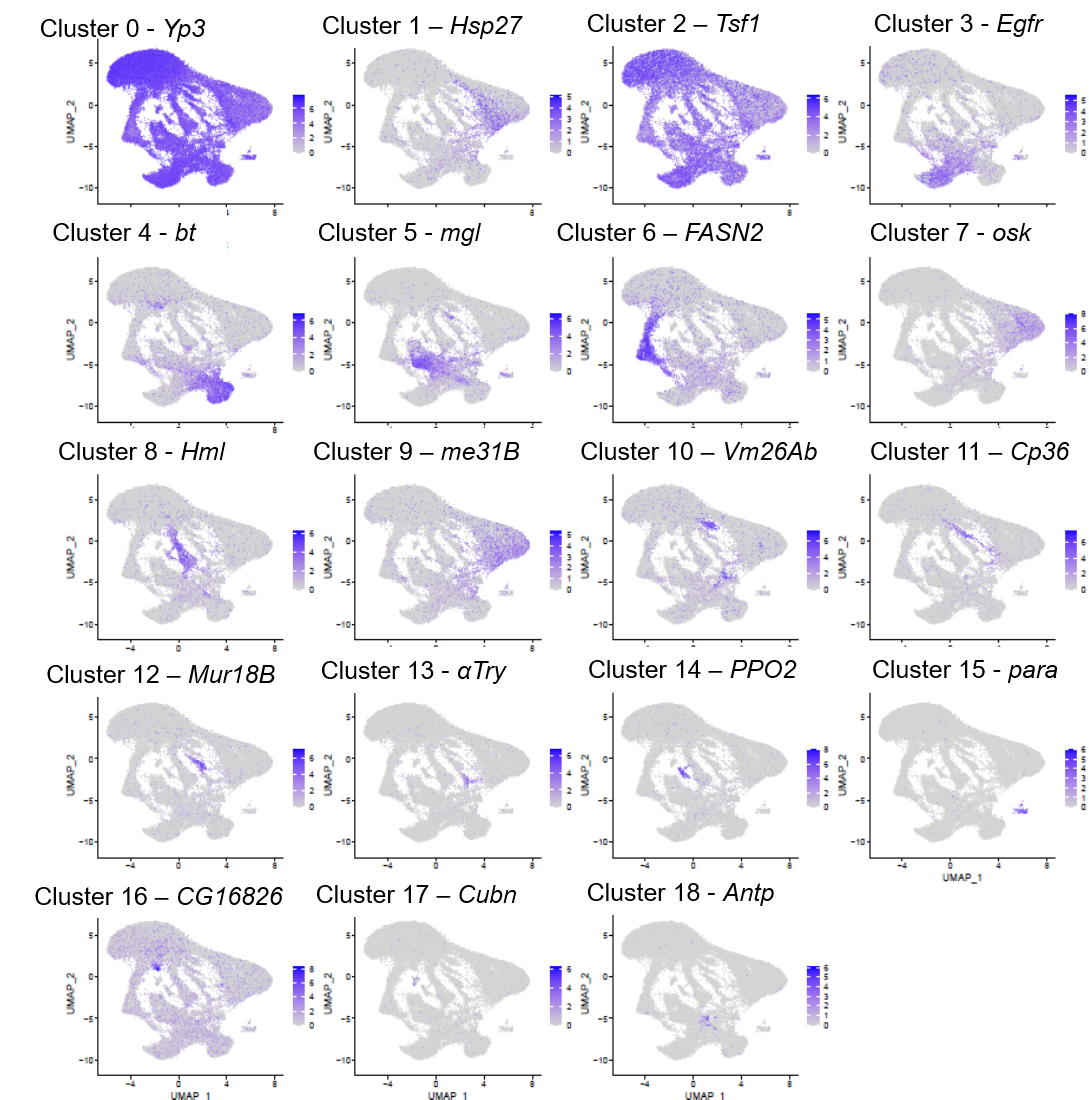

**Fig.S3: UMAP of expression of cluster-specific markers**.

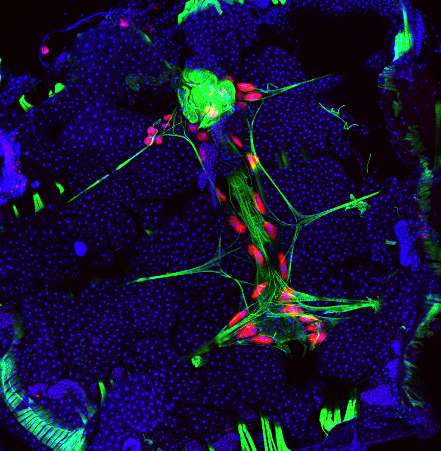

nACHR7α>mCherry

Fat Body

**Fig.S4.** **Spatial localization of Cluster 3**.

Expression of the fluorescent reporter mCherry under the control a nARCH*7a* driver labels putative Cluster 3 cells. The morphological and spatial profile of the cells expressing mCherry matched with that of pericardial cells (18,19). Nuclei are labelled with DAPI (blue) and actin is labeled with FITC-phalloidin (green).

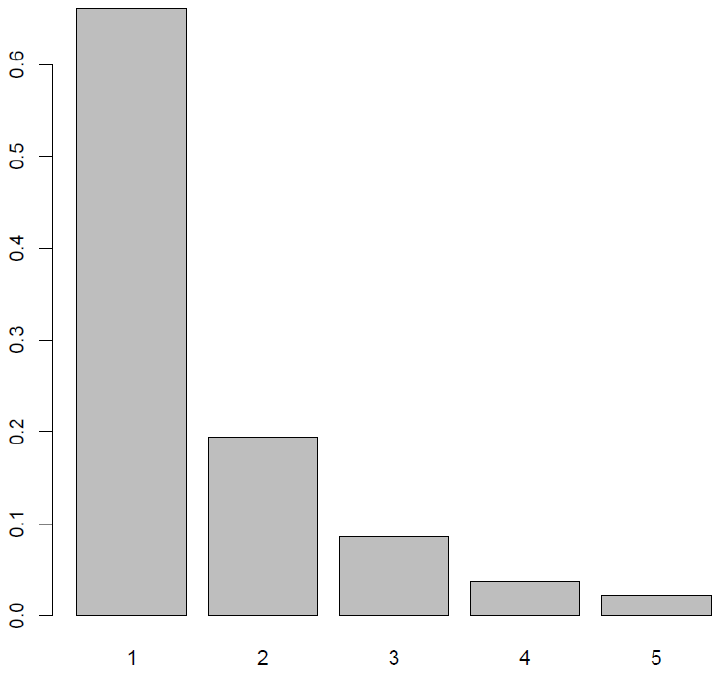

Proportion of differentially expressed genes

Number of Clusters

Genes differentially expressed between VU-MU

A.

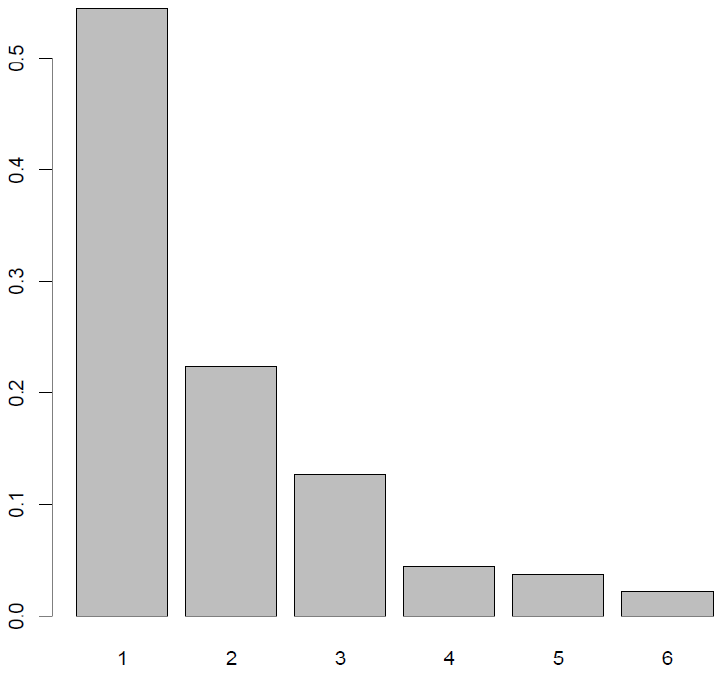

Number of Clusters

Genes differentially expressed between VU-VI

B.

Genes differentially expressed between MU-MI

Number of Clusters

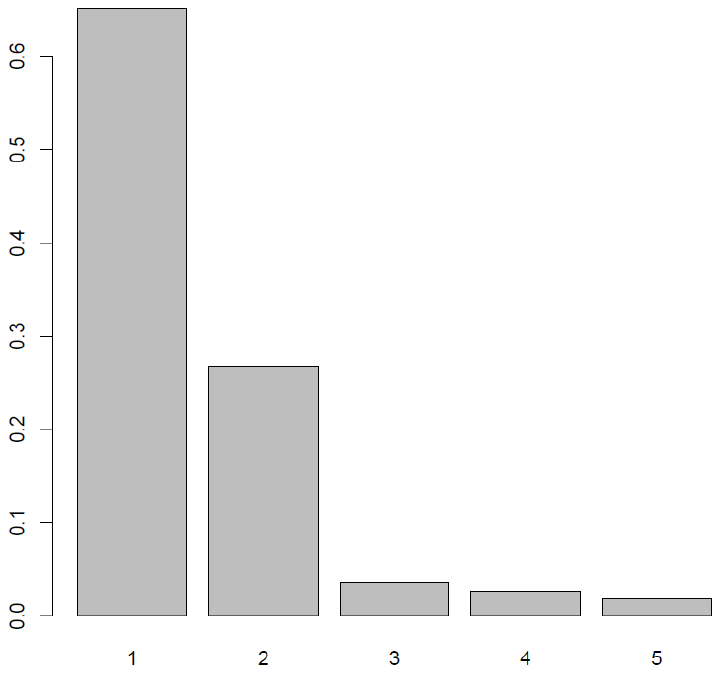

C.

**Fig.S5. Frequency distribution of genes that are differentially expressed between treatments across six fat body clusters**.

Proportion of differentially expressed genes

(A) Proportion of differentially expressed genes shared between clusters upon mating (Virgin Uninfected (VU) vs Mated Uninfected (MU)). (B) Proportion of differentially expressed genes upon infection in virgin females (Virgin Uninfected (VU) vs Virgin Infected (VI)). (C) Proportion of differentially expressed genes upon infection in mated females (Mated Uninfected (MU) vs Mated Infected (MI)).

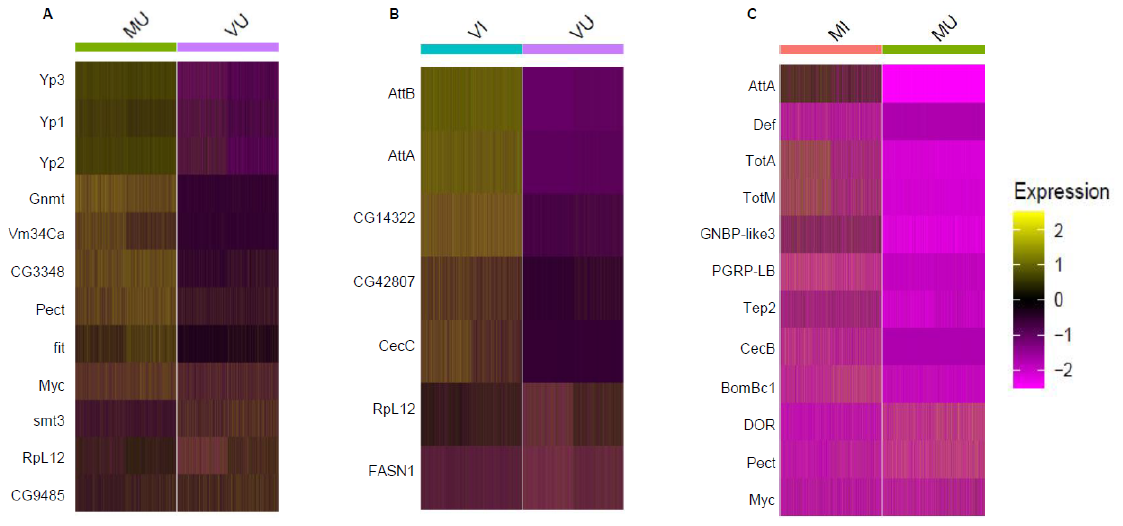

**Fig.S6. Heatmaps of top differentially expressed genes.**

(A) Upon mating (Virgin Uninfected (VU) vs Mated Uninfected (MU)), (B) Upon infection in Virgins (Virgin Uninfected (VU) vs Virgin Infected (VI)), and (C) Upon infection in Mated females (Mated Uninfected (MU) vs Mated Infected (MI)).

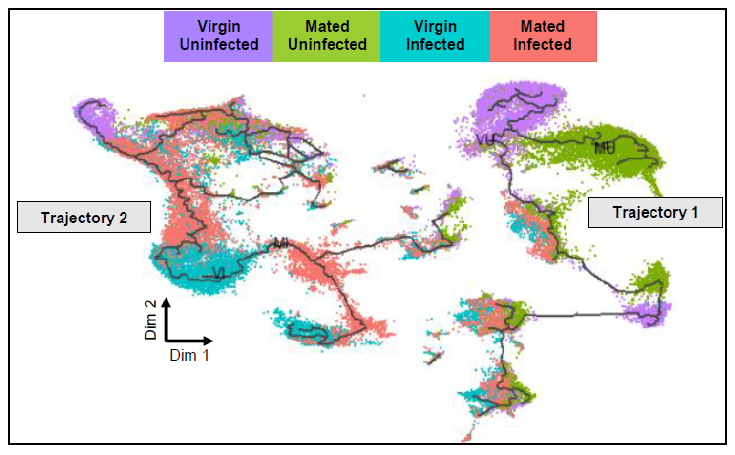

**Fig.S7. Trajectory analysis profiling of Drosophila fat body tissue**.

Monocle-based trajectory analysis separated nuclei into two dis-jointed trajectories (Trajectory 1 and Trajectory 2) primarily on the basis of their infection status. This indicates that nuclei from infected versus uninfected samples have dramatically different expression profiles Different colors represent nuclei from four different treatments (Virgin Uninfected, Mated Uninfected, Virgin Infected, and Mated Infected).

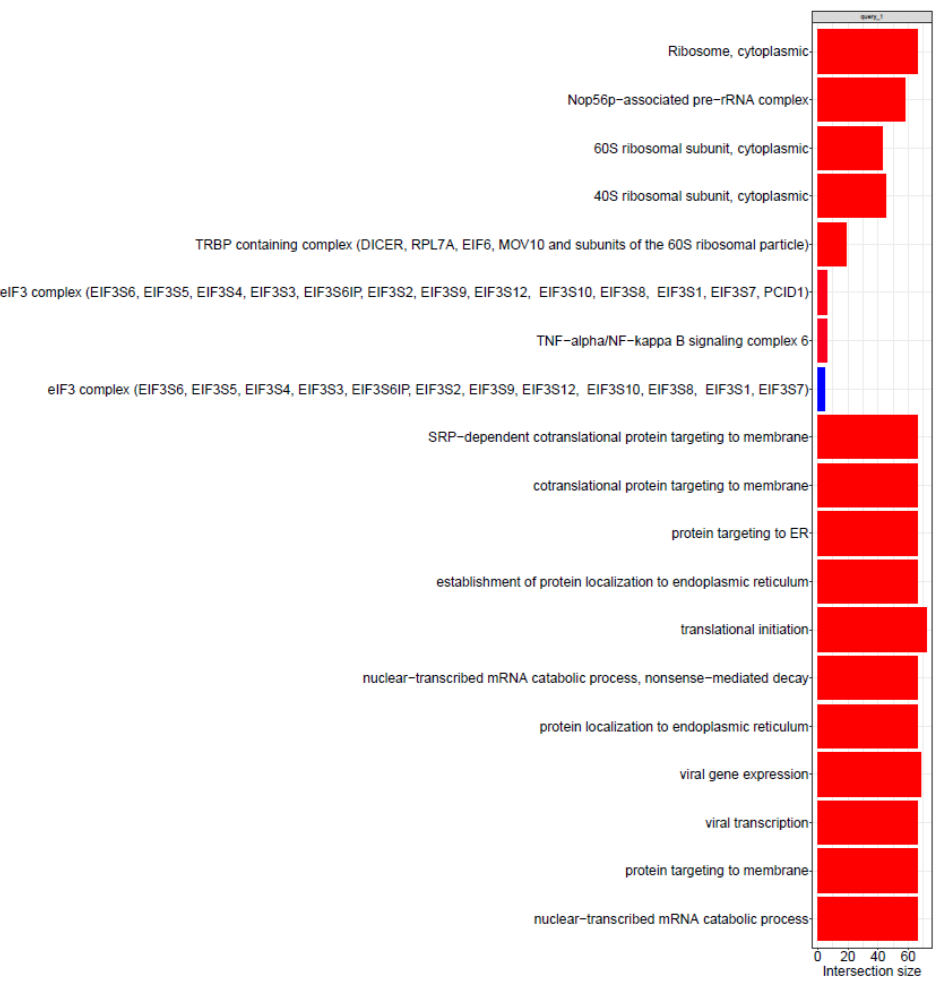

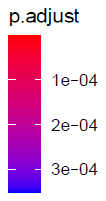

**Fig.S8. Enrichment plot of genes expressed in Module 13 obtained in pseudotime analysis**. Bar plot shows 20 most significantly enriched GO terms (p-adj). Enrichment was performed using g: profiler. P-adj: p-value for significant enrichment of GO terms after correction for multiple testing. Intersection size refers to the number of genes in the query that are annotated to each GO term. GO term analysis revealed enrichment for ribosome function and translation initiation in Module 13. Genes expressed in Module 13 showed lower gene expression score for Mated Uninfected compared to Virgin Uninfected (Fig. 2C).

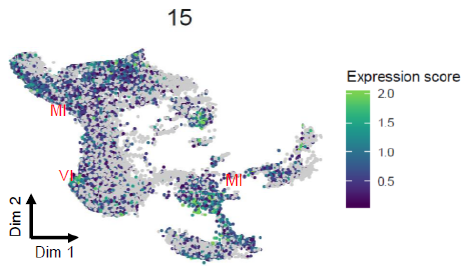

A.

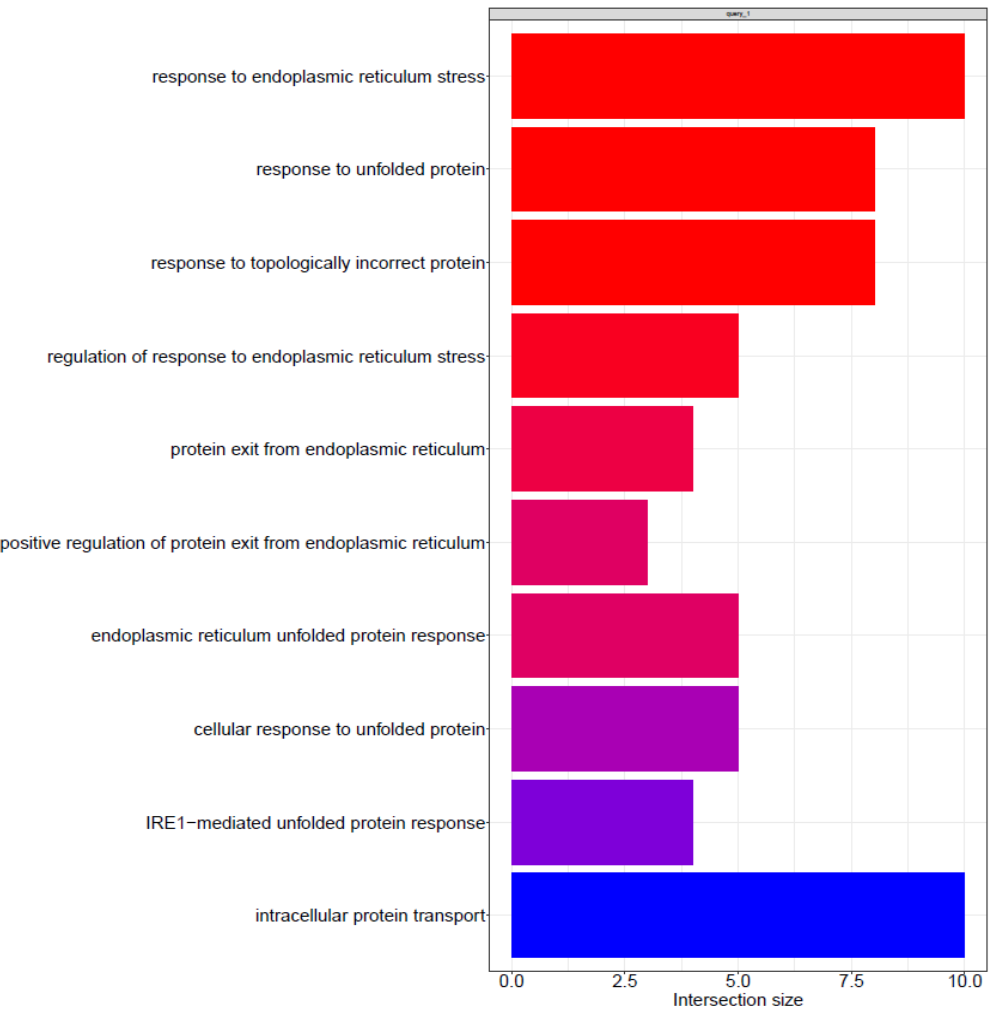

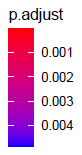

B.

**Fig.S9. Analysis of Module 15 (Trajectory 2) showing ER stress**.

(A) UMAP of Module 15 (Trajectory 2) showing high aggregate gene expression scores of genes present in Module 15 (Table S) for a subset of Mated Infected (MI) nuclei. Gradient of color represents the aggregate expression score with light color indicating higher expression score. Each dot represents a single nucleus. (B) Bar plot shows 10 most significantly enriched GO terms (p-adj) for Module 15. P-adj: p-value for significant enrichment of GO terms after correction for multiple testing. Intersection size refers to the number of genes in the query that are annotated to each GO term. GO term analysis showed enrichment for endoplasmic reticulum stress in Module 15.

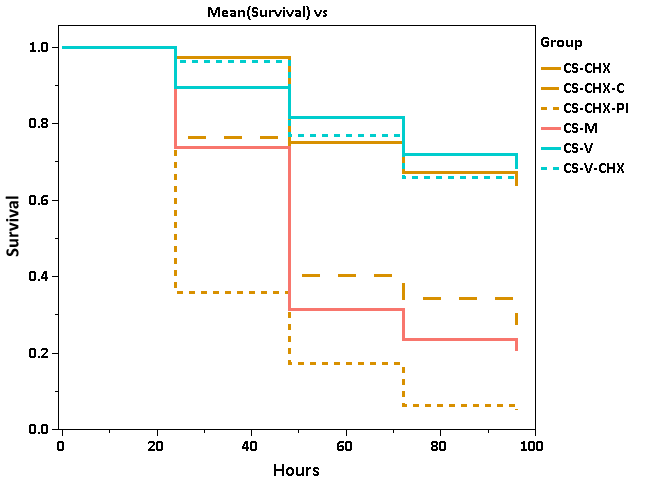

p < 0.0001

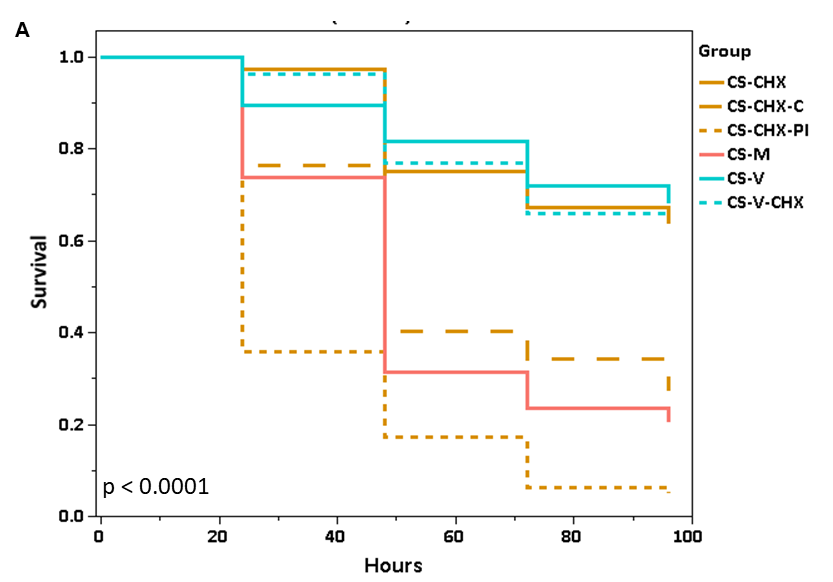

CHX

CHX - C

CHX - PI

MI

VI

CHX - VI

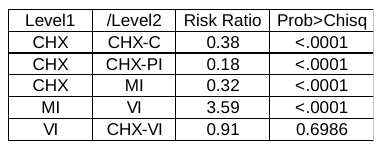

**Fig.S10. Survivorship analysis of post-infection mortality.**

Cox proportional hazard analysis showed that mated flies fed with Cycloheximide (CHX) **c**ontinuously (CHX-C) or only post-infection (CHX-PI) had significantly higher mortality compared to flies which were fed CHX only at the time of mating and before infection (CHX). Survivorship of virgin females fed with CHX (CHX-V) was not different from virgin females not fed CHX. Higher mortality of CHX-C and CHX-PI demonstrates that CHX does not provide any direct protection against infection with *Providencia rettgeri*. The CHX-C and CHX-PI treatments clearly demonstrate the necessity to synthesize proteins post-infection. (n = 35-40 flies per treatment per replicate, three replicates). p-value in the figure is the combined p-value of Cox proportional hazards analysis.

Table shows summary of results from risk ratios calculated between different treatments (Levels). Risk ratio is the ratio of hazard rates between two levels indicated as Level 1 and Level 2. Risk ratio < 1 shows lower risk to infection for level 1 and risk ratio > 1 shows higher risk to infection for level 1 – when compared to level 2.

| **Cluster#** | **Potential tissue** | **Marker genes** |
| --- | --- | --- |
| 0 | Fat Body Cells | *yp3, yp1* |
| 1 | Fat Body Cells | *Hsp27, dhd* |
| 2 | Fat Body Cells | *CG31326, Nep6* |
| 3 | Unknown | *Pde9, Ubx* |
| 4 | Muscle Cells | *bt, sls* |
| 5 | Proliferative Fat Body Cells | *mgl, CG14661* |
| 6 | Oenocytes | *FASN2, CG7910* |
| 7 | Reproductive provisioning fat body cells | *osk,dhd* |
| 8 | Hemocytes | *Hml,Ten-m* |
| 9 | Uncharacterized | *me31B, Hsp27* |
| 10 | Structural fat body cells | *Vm26Ab, Vm26Aa* |
| 11 | Chorion producing fat body cells | *Cp36, Cp38* |
| 12 | Catabolic fat body cells | *Mur18b, CG14292* |
| 13 | Putative digestive cells | *alphaTry, Jon65Aiv* |
| 14 | Crystal cells | *PPO1, PPO2* |
| 15 | Neuronal | *Para, Rdl* |
| 16 | Stress response fat body cells | *whe, CG16826* |
| 17 | Nephrocytes | *Cubn, CG42255* |
| 18 | Tracheal cells | *Antp, trol* |

Table S1: A summary of potential tissue type and markers representing each cluster. 19 clusters were identified from the entire dataset.

Table S7: A summary of the number of genes that are differentially expressed between treatments across 19 clusters. **VU-MU**: Number of genes differentially expressed upon mating (Virgin Uninfected (VU) vs Mated Uninfected (MU)), **VU-VI**: Number of genes differentially expressed upon infection in Virgins (Virgin Uninfected (VU) vs Virgin Infected (VI)), **MU-MI**: Number of genes differentially expressed upon infection in Mated (Mated Uninfected (MU) vs Mated Infected (MI)).

|  | Differentially expressed genes compared between | | | | | |
| --- | --- | --- | --- | --- | --- | --- |
|  | VU-MU | | VU-VI | | MU-MI | |
| Cluster# | Upregulated | Downregulated | Upregulated | Downregulated | Upregulated | Downregulated |
| 0 | 26 | 1 | 45 | 1 | 26 | 10 |
| 1 | 82 | 12 | 41 | 0 | 8 | 23 |
| 2 | 58 | 26 | 84 | 8 | 46 | 25 |
| 3 | 82 | 52 | 88 | 84 | 56 | 62 |
| 4 | 51 | 3 | 41 | 5 | 14 | 7 |
| 5 | 51 | 1 | 33 | 2 | 13 | 6 |
| 6 | 26 | 2 | 25 | 0 | 3 | 6 |
| 7 | 20 | 5 | 25 | 0 | 3 | 2 |
| 8 | 33 | 1 | 48 | 4 | 14 | 11 |
| 9 | 12 | 15 | 2 | 0 | 0 | 1 |
| 10 | 6 | 3 | 12 | 0 | 4 | 1 |
| 11 | 3 | 0 | 2 | 0 | 1 | 0 |
| 12 | 3 | 0 | 7 | 0 | 0 | 0 |
| 13 | 0 | 0 | 2 | 0 | 0 | 0 |
| 14 | 2 | 0 | 6 | 0 | 2 | 0 |
| 15 | 0 | 0 | 2 | 0 | 0 | 0 |
| 16 | 0 | 0 | 0 | 0 | 0 | 1 |
| 17 | 1 | 0 | 0 | 0 | 0 | 0 |
| 18 | 0 | 0 | 0 | 0 | 0 | 0 |

Table S11: (A) Effect of treatment (VU, VI, MU, and MI) on puromycin incorporation (relative fluorescence) measured using Western blotting. Summary of results of one-way ANOVA with Treatment using levels VU, VI, MU, and MI. Posthoc comparisons using Tukey’s HSD shows significantly different puromycin incorporation between VI and MI (Fig.4B). (B) Summary of results of pairwise contrast between VI and MI.

A.

| Source | DF | Sum of Squares | Mean Square | F Ratio | Prob > F |
| --- | --- | --- | --- | --- | --- |
| Treatment | 3 | 5.677495 | 1.8925 | 4.26 | 0.0216 |
| Error | 16 | 7.108 | 0.44425 |  |  |
| C. Total | 19 | 12.785495 |  |  |  |

B.

| Effect | SS | NumDF | DenDF | F Ratio | Prob > F |
| --- | --- | --- | --- | --- | --- |
| Genotype | 4.80249 | 1 | 16 | 10.8103 | 0.0046 |

Table S12: *t-test* results comparing effect of cycloheximide treatment on puromycin incorporation. MI represents Mated Infected females not treated with cycloheximide and CHX represents females treated with CHX (See Methods).

| Treatment | | | | | |  |
| --- | --- | --- | --- | --- | --- | --- |
| MI | | | CHX | | | *t-test* |
| N | Mean | SD | N | Mean | SD | 3.6** |
| 5 | 1.02 | 0.89 | 5 | 1.59 | 0.61 |  |
| ***p* = 0.02 | |  |  |  |  |  |

**The following tables have been provided as separate files (.xlsx)**

**Table S2. Differentially expressed genes upon mating**. padj = P-value adjusted for multiple testing. fc = log fold-change between the two treatments. Pct. 1 = percentage of cells in Mated Uninfected (MU) expressing the gene. Pct. 2 = percentage of cells in Virgin Uninfected (VU) expressing the gene.

**Table S3. Enrichment analysis** of six identified fat body clusters 0,1,2,5,7, and 10 using genes differentially expressed between Mated uninfected (MU) and Virgin Uninfected (VU). adjusted_p_value **=** P-value adjusted for multiple testing.

**Table S4. Differentially expressed genes upon infection in virgins**. padj = P-value adjusted for multiple testing. fc = log fold-change between the two treatments. Pct. 1 = percentage of cells in Virgin Infected (VI) expressing the gene. Pct. 2 = percentage of cells in Virgin Uninfected (VU) expressing the gene.

**Table S5. Differentially expressed genes upon infection in mated females**. padj = P-value adjusted for multiple testing. fc = log fold-change between the two treatments. Pct. 1 = percentage of cells in Mated Infected (MI) expressing the gene. Pct. 2 = percentage of cells in Mated Uninfected (MU) expressing the gene. adjusted_p_value = P-value adjusted for multiple testing.

**Table S6**. **VI-VU**: **Enrichment analysis** of six identified fat body clusters 0,1,2,5,7, and 10 using genes differentially expressed between Virgin Uninfected (VU) and Virgin Infected (VI).

**MI-MU**: **Enrichment analysis** of six identified fat body clusters 0,1,2,5,7, and 10 using genes differentially expressed between Mated Uninfected (MU) and Mated Infected (MI). adjusted_p_value = P-value adjusted for multiple testing.

**Table S8. Genes expressed in module 13 of partition 1.** Genes expressed in this module show low aggregate gene expression score in Mated Uninfected.

**Table S9. Genes expressed in module 16 of partition 2.** Genes expressed in this module show low aggregate gene expression score in Mated Infected.

**Table S10. Genes expressed in module 15 of partition 2.** Genes expressed in this module show high aggregate gene expression score in Mated Infected.

**Table S13: Marker genes for each cluster** (Clusters 0 to 18) obtained using function 'FindConservedMarkers' in Seurat. Avg_logFC: average log fold change between cluster of interest and all other clusters. minimump_p_val: combined p-value for all the four treatments (Virgin Uninfected. Virgin Infected, Mated Uninfected, Mated Infected)

**Table S14: Enrichment analysis of marker genes for each cluster (Clusters 0 to 18).**
